# Image-based consensus molecular subtype classification (imCMS) of colorectal cancer using deep learning

**DOI:** 10.1101/645143

**Authors:** Korsuk Sirinukunwattana, Enric Domingo, Susan Richman, Keara L Redmond, Andrew Blake, Clare Verrill, Simon J Leedham, Aikaterini Chatzipli, Claire Hardy, Celina Whalley, Chieh-Hsi Wu, Andrew D Beggs, Ultan McDermott, Philip Dunne, Angela A Meade, Steven M Walker, Graeme I Murray, Leslie M Samuel, Matthew Seymour, Ian Tomlinson, Philip Quirke, Tim Maughan, Jens Rittscher, Viktor H Koelzer, on behalf of S:CORT consortium

## Abstract

Image analysis is a cost-effective tool to associate complex features of tissue organisation with molecular and outcome data. Here we predict consensus molecular subtypes (CMS) of colorectal cancer (CRC) from standard H&E sections using deep learning. Domain adversarial training of a neural classification network was performed using 1,553 tissue sections with comprehensive multi-omic data from three independent datasets. Image-based consensus molecular subtyping (imCMS) accurately classified CRC whole-slide images and preoperative biopsies, spatially resolved intratumoural heterogeneity and provided accurate secondary calls with higher discriminatory power than bioinformatic prediction. In all three cohorts imCMS established sensible classification in CMS unclassified samples, reproduced expected correlations with (epi)genomic alterations and effectively stratified patients into prognostic subgroups. Leveraging artificial intelligence for the development of novel biomarkers extracted from histological slides with molecular and biological interpretability has remarkable potential for clinical translation.

## INTRODUCTION

Colorectal cancer (CRC) is a disease with heterogeneous molecular subtypes, variable clinical course and prognosis (*1*). An increasing understanding of CRC biology has led to the development of targeted treatments directed against key pro-oncogenic signalling pathways, but these treatments are only effective in a small proportion of patients (*2, 3*). Molecular stratification of CRC patients is essential to form homogenised subgroups for personalised treatment and prognosis (*4*). Next generation sequencing (NGS) technologies enable the multi-omic profiling of malignant tumours but impact on clinical practice has been limited. This is due to high costs, difficulty in the standardisation of pre-analytical procedures, requirements for data storage and bioinformatics expertise (*5, 6*). In contrast, histopathology slides are inexpensive to produce and principal stains such as haematoxylin and eosin (H&E) are firmly established in the pathology lab.

The application of traditional image analysis to histopathology facilitates the quantitative assessment of tissue architecture, cell distribution, and cellular morphology by light microscopy to generate feature libraries of unprecedented resolution and detail (*7*). More recently, deep learning is used to capture morphological differences with a precision that exceeds human performance. Coudray et al utilise this approach to detect targetable oncogenic driver mutations in lung cancer using deep neural classification networks (*8*). By combining an image-based analysis with molecular characterisation, it becomes feasible to identify novel genotype-phenotype correlations. For the first time it is now possible to characterise complex multi-scale morphological traits as well as genomic alterations at scale. Given that H&E processing allows analysis of large tissue sections at low cost and with short turn around without the need to modify existing clinical workflows, the discovery of morpho-molecular correlations holds the promise of revolutionising patient stratification in clinical practice (*9*). Image-based methods are suitable for prioritisation of certain patient samples for additional molecular testing and for provision of additional guidance for the selection of tissue blocks. Ultimately, the biological interpretability of genomic alterations could revolutionise the development of new biomarkers.

In CRC, it is well known that tumour morphology, growth pattern and architecture hold important clues to differentiating biological subtypes with clinical impact (*10*). The composition of the tumour microenvironment is a key component determining the tumour progression and therapy response (*11, 12*). Tumour and non-tumour tissue contribute to image information on the histological slide and to the consensus molecular classification (CMS) of CRC at the transcriptional level (*13*). The CMS classification distinguishes four groups of CRC with distinct clinical behaviour and biological interpretability. These include CMS1 (14%; microsatellite instability immune, favourable prognosis), CMS2 (37%, canonical, epithelial gene expression profile, WNT and MYC signalling activation, intermediate prognosis), CMS3 (13%, epithelial profile with evident metabolic dysregulation, intermediate prognosis), and CMS4 (23%, mesenchymal, prominent transforming growth factor-β activation, poor prognosis) (*1, 13*).

CMS subgrouping shows a robust association with targetable alterations and may have potential to guide treatment allocation in clinical practice (*1, 13*). However, clinical implementation of the CMS classification has been held back by the considerable costs of RNA sequencing, the inability to bioinformatically obtain confident CMS calls from single samples, intratumoural heterogeneity, high levels of unclassified calls on biopsies and an unclear performance on FFPE material (*13–15*). Here, we derive a novel image-based CMS (imCMS) classification from H&E-stained tissue sections sourced from the Medical Research Council (MRC) and Cancer Research UK (CRUK) Stratification in COloRecTal cancer (S:CORT) program and The Cancer Genome Atlas (TCGA). We demonstrate the existence of distinct image phenotypes of CRC that reproducibly associate with CMS transcriptional classification, key oncogenic driver mutations and prognosis. Automatic, high-fidelity classification of three independent clinical cohorts including pre-operative biopsies underlines the applicability of this approach to heterogeneous sample sets and relevant clinical settings. We provide insight into classification calls for samples with considerable intratumoural heterogeneity and provide accurate secondary calls with higher discriminatory power than bioinformatic prediction. In all three cohorts, imCMS successfully classified CRC samples that were previously considered to have unknown biological and clinical behaviour and failed transcriptional classification. imCMS classification is standardised, inexpensive and could be carried out in a tele-pathology setting on routinely available H&E sections. This resolves key issues in the translation of transcriptional classification of CRC into clinical practice and has the potential to increase availability of molecular stratification in low resource settings.

## MATERIALS AND METHODS

### Study design

This tudy was designed in accordance with the REMARK guidelines. The study design, cohorts and aims are outlined in [**Figure 1**].

**Figure 1:**
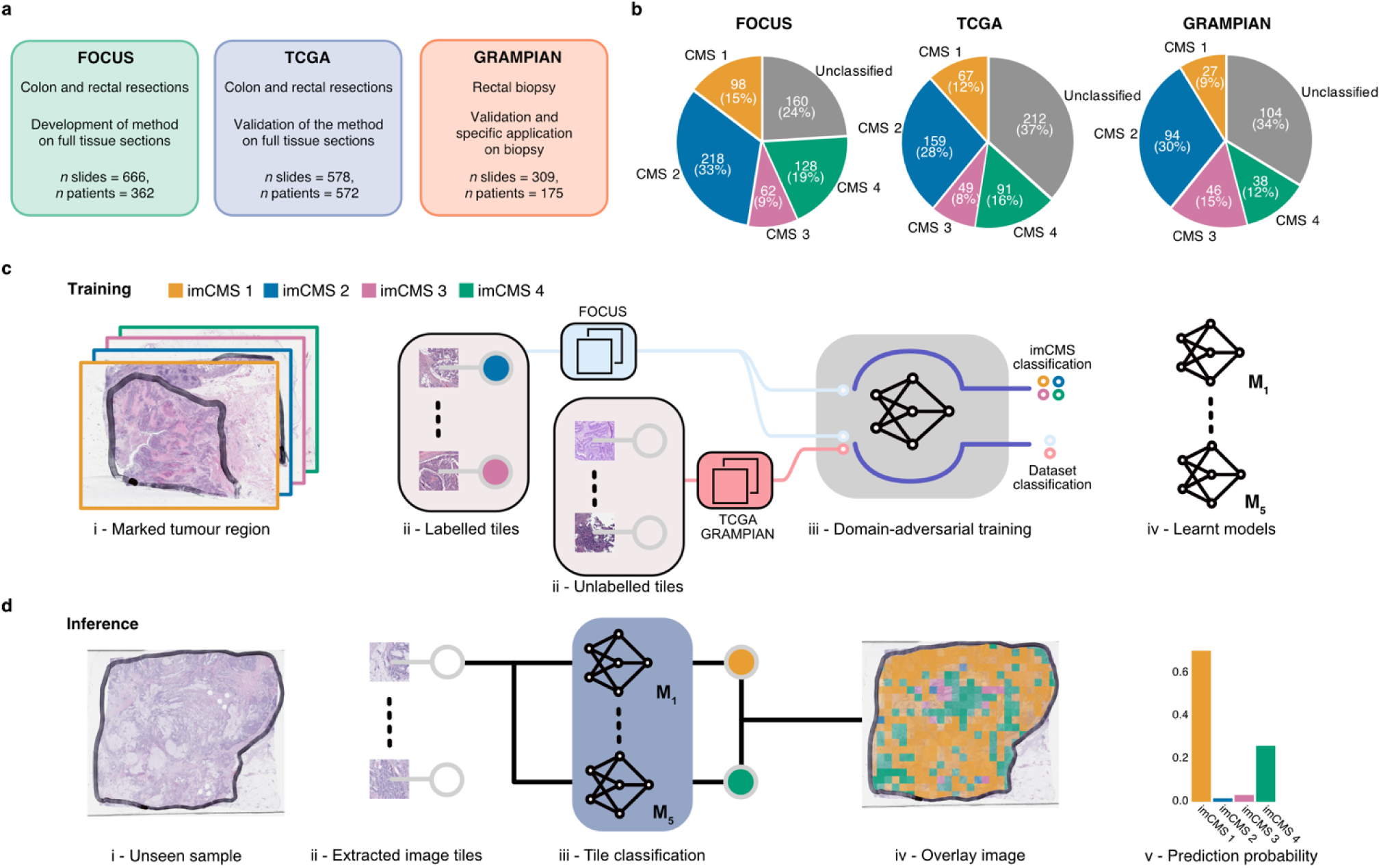
Data, study design, and imCMS classification framework. Three independent datasets (FOCUS, TCGA, and GRAMPIAN) were used in this study. **(A)** The distribution of the samples stratified by the CMS calls in each dataset. **(B)** The FOCUS dataset was primarily used for learning the imCMS discriminative model while the TCGA and GRAMPIAN datasets were used for testing. **(C)** Training of the imCMS discriminative model based on the domain-adversarial approach. Image tiles were extracted from annotated tumour regions. Tiles from the FOCUS cohort were categorised by CMS class of the original slide and were used to train the model to predict the imCMS classes on unseen datasets. Tiles from the TCGA and GRAMPIAN cohorts were unlabelled and were used together with those from the FOCUS cohort in the cohort (domain) prediction. Domain- adversarial training forced the cohort classifier to perform poorly which in turn encouraged the model to learn indiscriminative features across datasets. Five distinct models were produced. **(D)** At the inference time, the ensemble of the learnt models predicts the imCMS class for each of the image tiles extracted from annotated tumor regions of a slide. A slide is assigned to the imCMS class with the maximum prediction score (i.e. highest number of tiles in the slide).

### Patients

#### Cohort 1: FOCUS (Retrospective cohort, S:CORT)

As part of the Stratification in COloRecTal cancer (S:CORT) program, 385 patients with available formalin-fixed paraffin embedded (FFPE) blocks of the primary CRC were selected from the MRC FOCUS randomised clinical trial (RCT) that tested different strategies of sequential and combination chemotherapy for patients with advanced CRC (*30*). Serial sections were cut from one representative block for H&E staining followed by four unstained sections for RNA extraction, a second H&E and eight unstained sections for DNA extraction for a total of 741 slides. H&E slides were re-reviewed by expert gastrointestinal pathologists and tumour tissue was annotated and used to guide RNA and DNA extractions from the first and second H&E respectively. RNA expression microarrays (Xcel array, Affymetrix), DNA target capture (SureSelect, Agilent) followed by NGS sequencing (Illumina) and DNA methylation arrays (EPIC arrays, Illumina) were applied in this order. All H&E slides were scanned at high resolution on an Aperio scanner at a total magnification of 200X. Digital slides were re-reviewed and tumour annotations were traced to generate region annotations for machine learning classification. Clinical data was retrieved from the trial database. Pathological TNM-stage and sidedness were extracted from pathological reports. Patients with synchronous disease were considered to be stage IV. 34 slides with technical failure of the staining or scanning procedure were excluded from further analysis. 41 slides had no available RNA expression for CMS classification for a final set of 666 slides (n=362 cases). Clinical and molecular data is summarised in [**Table S1**] and [**Figure 1A-B**].

#### Cohort 2: TCGA (colon and rectal adenocarcinomas)

A total of 623 digital slides from 614 cases of colon and rectal adenocarcinoma with available FFPE samples were downloaded from the TCGA Data Portal (data accessed on August 2nd, 2018). All digital slides were re-reviewed and tumour tissue was annotated. A total of 45 slides were excluded based on quality control criteria. Clinical data was obtained from Liu et al (*31*) while somatic mutations and gene level expression data were downloaded with the R package TCGAbiolinks (*32*) on November 7th, 2018. Mutations from Varscan and Mutect were combined and calls for driver mutations were computed for relevant genes (all truncating mutations for *APC*; missense mutations for *KRAS* in codons 12, 13, 19, 22, 59, 61, 68, 117 and 146; V600E for *BRAF;* all missense and truncating mutations for *TP53*). The final number of slides for imCMS classification was 578 (n=572 patients) [**Table S1**] and [**Figure 1A-B**].

#### Cohort 3: GRAMPIAN (Retrospective cohort, S:CORT)

A total of 323 slides from 183 pre-treatment biopsy FFPE blocks from rectal cancer patients of the neoadjuvant setting were available for this study as part of the S:CORT program. All patients received pre-operative chemoradiotherapy followed by surgical resection. Slides and molecular profiling were processed as described for cohort 1 (FOCUS) but using 5 to 9 sections for RNA extraction and 9 for DNA. Pre-operative staging was derived from MRI scans. A total of 14 slides were excluded based on quality control criteria for a final set of 309 slides (n = 175 cases). Clinical and molecular data is summarised in [**Table S1**] and [**Figure 1A-B**].

### Assay methods

#### CMS calls

RNA microarray data was pre-processed and normalised using robust multi-array analysis with the R package affy (*33*) and probes collapsed by mean. CMS calls in all three cohorts were derived with the R package CMSclassifier (*13*) by random forest (RF) with the default posterior probability of 0.5. RF CMS classification of FFPE samples from the FOCUS and GRAMPIAN cohorts led to an increased frequency of unclassified samples as compared to the TCGA datasets derived from fresh frozen material. In order to derive calls with comparable frequencies, we therefore computed single sample predictor calls (R package CMSclassifier) after row-centring the expression data (*13*). Final CMS calls were generated when there was a match between both methods (RF and single sample predictor without applying any cut-off). There were 186 TCGA cases (n=191 slides) with discrepancies among our CMS calls and the calls originally reported by Guinney et al (*13*). These discrepant calls are most likely the result of the application of a clustering method that is strongly cohort-dependent in our analysis based on TCGA samples only and the original report combining thousands of samples from several selected cohorts. Due to lack of clear evidence of the ground truth CMS status, samples with classification discrepancies were labelled as unclassified.

Secondary CMS calls from RNA in classified samples were computed by RF using the second highest call with posterior probability above 0.3. The primary call was matched if no different CMS subtype was found. For unclassified samples, the first highest call above 0.3 was used, leaving the sample as unclassified if no subtype met this requirement. All these analyses were performed with R version 3.5.1 (*34*).

#### CIMP classification

Methylation array raw data from S:CORT cohorts 1 and 3 was processed with the R-package ChAMP (*35*). CIMP classification was generated by recursively partitioned mixture model as previously done in TCGA (*36*) and Guinney et al (*13*) with minor changes due to the higher number of probes. CIMP classification in TCGA according to Guinney et al was retrieved from Synapse (Synapse ID syn2623706).

### imCMS classification

#### Pre-processing of image data and exclusion criteria

For each of the three cohorts, digital slides were re-reviewed and invasive cancer regions were annotated by an expert gastrointestinal pathologist using the HALO^TM^ software v2.3.2089.52 (Indica Labs, Corrales, NM, USA). For each slide, the annotated tumour areas were divided into tiles of 512x512 pixels. To avoid white background regions which did not provide useful information for classification, we excluded tiles with less than 50% tissue area. Total tissue area and the number of tiles is shown in [**Figure S1**]. At 5x magnification, consecutive tiles were 50% overlapped in the FOCUS and TCGA cohorts (resections). To account for the small sample surface area of the tumour identified in the endoscopic biopsies of the GRAMPIAN cohort at 5x, tiles with a 75% overlap were used. At 20x, no overlap in FOCUS and TCGA and 50% overlap in GRAMPIAN were used.

#### imCMS classifier and the training procedure

We trained a neural network to classify a given image tile taken from the marked tumour area into one of the four CMS classes using supervised learning. Inception V3 (*37*) pretrained on the ImageNet dataset (*38*) was trained on samples taken from the FOCUS cohort [**Figure 1C**]. All instances in the training set were associated with corresponding molecular data. The class of each tile in the training set was matched to the overall RNA-based CMS call of the FOCUS slide. Tiles from unclassified slides were excluded. We trained 5 separate models with different subsets of the data in the manner akin to cross-validation. The data were split into 5 partitions while preserving the percentage of samples for each CMS class. For each model, 3 portions of the data were used for training, one for validation, and one for testing. The split was done at the patient level, meaning that no image tiles from the same patients would be used for training, validation, and testing at the same time. An inception V3 (*37*) model pretrained on the ImageNet dataset (*38*) was deployed. We minimised the cross-entropy loss of the model on our dataset via gradient backpropagation using Adam optimisation (*39*) with a learning rate of 0.0002 and a batch size of 32 for 100,000 iterations. To prevent the model from overfitting, the training image tiles were aggressively augmented using diverse optical and spatial transformations implemented in the imgaug library (*40*). To further avoid the class imbalance problem, we also sampled tiles according to the inverse of their class frequencies to guarantee that tiles from the minority classes such as CMS3 were sampled frequently in the training process. Finally, we selected the state of the model that yields the maximum macro-average AUC on the validation data. We implemented the entire imCMS classification framework using the deep learning Pytorch library (*41*). All statistical analyses were performed in R version 3.5.1 (*34*).

#### Testing the model on independent cohorts

On the TCGA and GRAMPIAN datasets, we applied 5 versions of the, producing 5 different classification results for each tile which were then averaged to obtain the final prediction. This is analogous to an ensemble of experts’ opinions (*27*). The prediction probability for each imCMS class was obtained from the proportion of the number of tiles assigned to that class, and the final imCMS call at the slide level was derived from the majority vote of tiles [**Figure 1D**]. No unclassified slides were used in the evaluation. The classification performance of the model is reported in [**Table 2**].

**Table 1:** Area under the curve (AUC) with 95% confidence intervals achieved by the imCMS classifier.

| CMS class | ***FOCUS***<br>n slides = 506<br>n patients = 276 |  | ***TCGA***<br>n slides = 366<br>n patients = 365 |  | ***GRAMPIAN***<br>n slides = 205<br>n patients = 114 |  |
| --- | --- | --- | --- | --- | --- | --- |
|  | 5x | 20x | 5x | 20x | 5x | 20x |
| **CMS1** | 0.85 (0.8,0.89) | 0.85 (0.81,0.89) | 0.8 (0.73,0.87) | 0.8 (0.75,0.88) | 0.73 (0.6,0.9) | 0.79 (0.72,0.87) |
| **CMS2** | 0.88 (0.86,0.91) | 0.86 (0.83,0.91) | 0.79 (0.74,0.83) | 0.79 (0.75,0.83) | 0.76 (0.69,0.83) | 0.76 (0.7,0.83) |
| **CMS3** | 0.92 (0.9,0.96) | 0.9 (0.85,0.94) | 0.77 (0.68,0.88) | 0.74 (0.65,0.82) | 0.81 (0.74,0.89) | 0.85 (0.78,0.92) |
| **CMS4** | 0.86 (0.83,0.9) | 0.85 (0.82,0.89) | 0.78 (0.72,0.86) | 0.77 (0.72,0.82) | 0.92 (0.87,0.99) | 0.92 (0.88,1) |
| **Macro-average** | 0.88 (0.86,0.9) | 0.87 (0.84,0.89) | 0.79 (0.75,0.83) | 0.78 (0.74,0.81) | 0.81 (0.76,0.85) | 0.83 (0.79,0.88) |
| **Micro-average** | 0.89 (0.88,0.91) | 0.88 (0.86,0.91) | 0.77 (0.74,0.8) | 0.77 (0.73,0.81) | 0.83 (0.8,0.87) | 0.85 (0.81,0.89) |

**Table 2:**
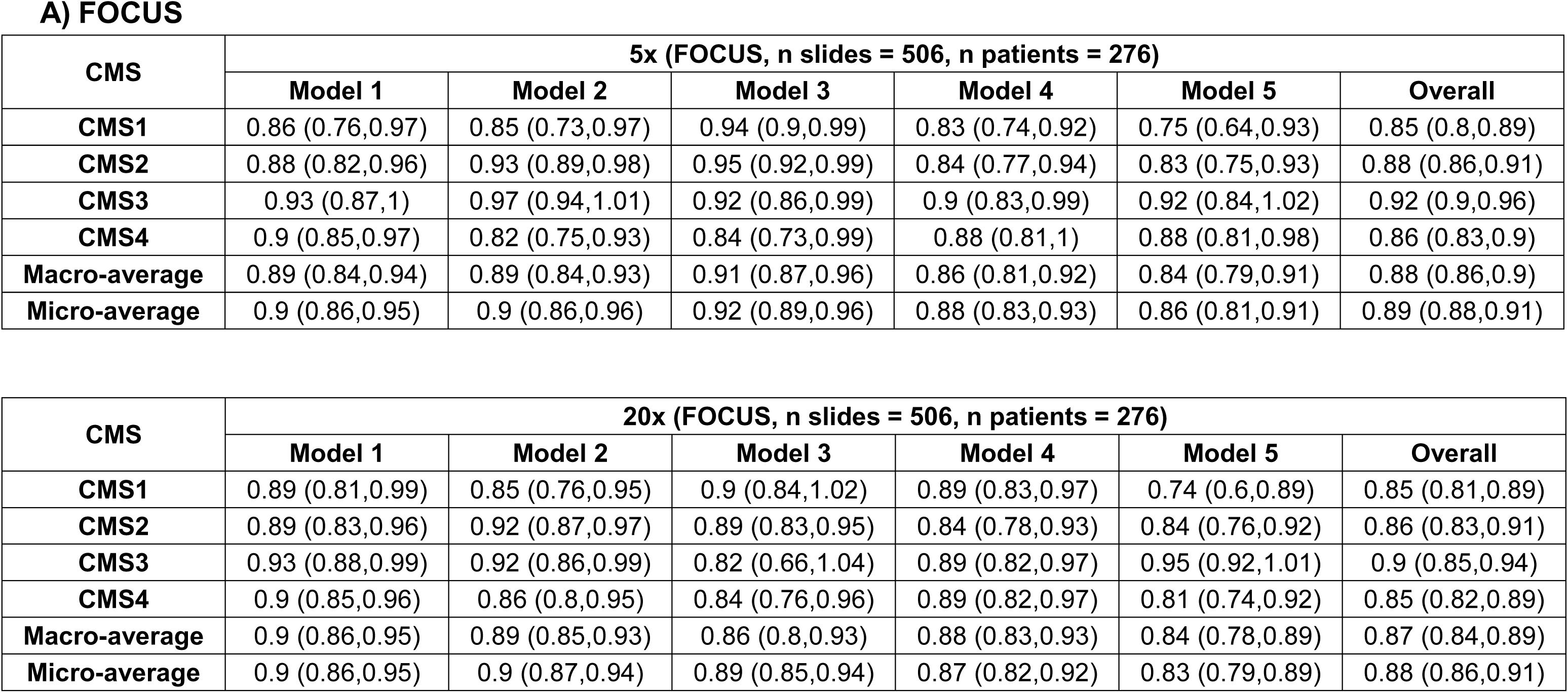
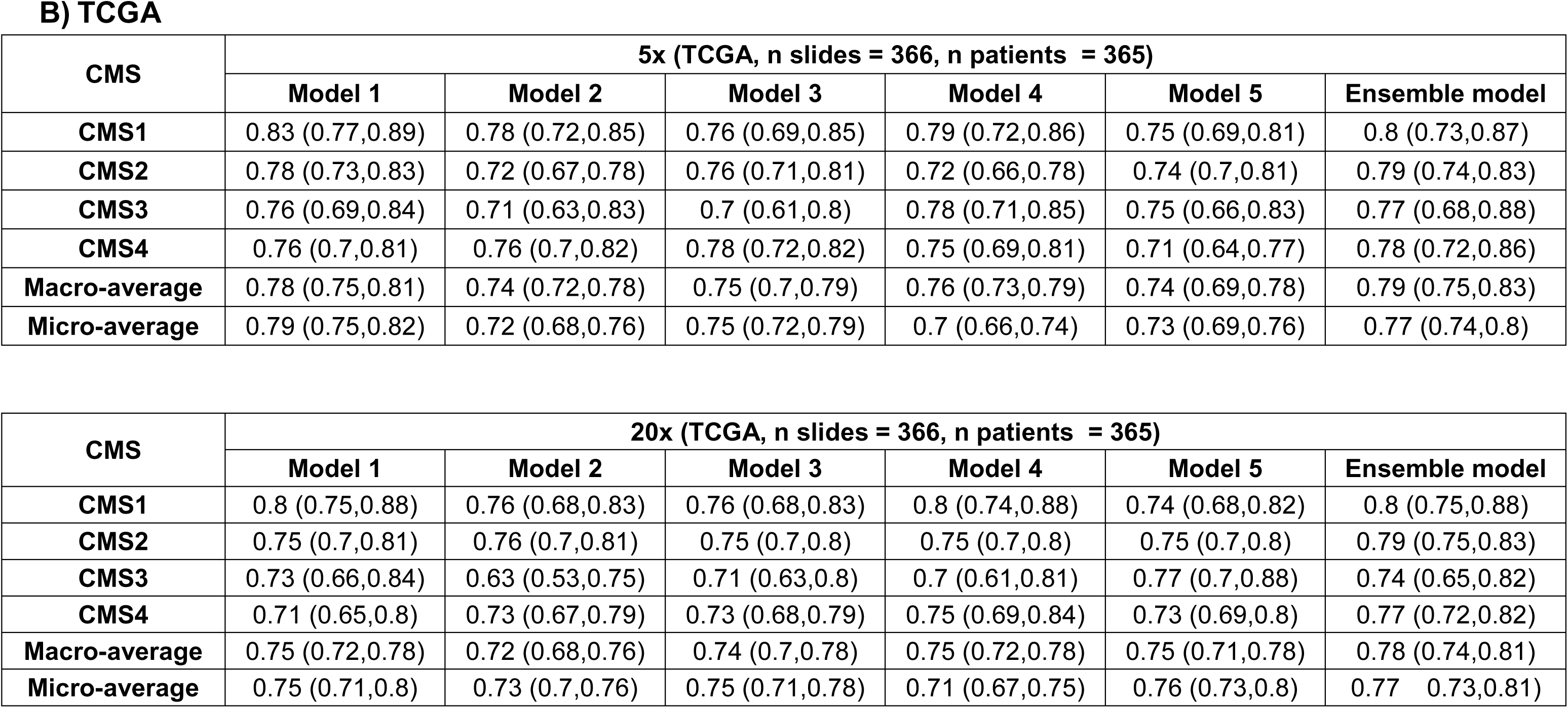
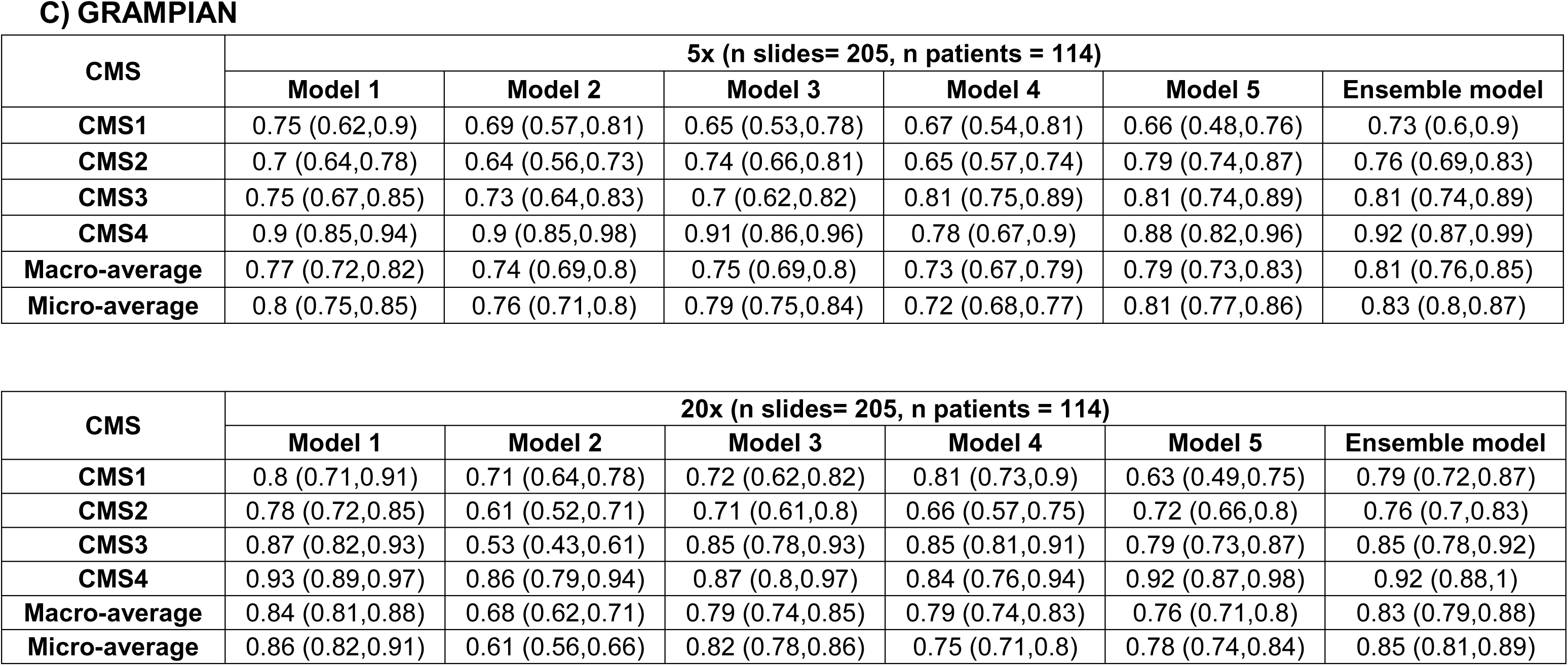
Area under the curve (AUC) with 95% confidence intervals achieved by the imCMS classifier.

#### Domain adversarial training for better generalisation

To prevent the learning of dataset-dependent features that would limit the general applicability of the model we leveraged domain-adversarial training (*26*). Here the model was augmented with an additional classifier for predicting whether image tiles were drawn from training (FOCUS) or external cohorts (TCGA and GRAMPIAN) [**Figure 1C**]. We forced this classifier to perform poorly to encourage the model to learn features which are dataset-independent. To train the domain-adversarial classifier, all image tiles from the FOCUS cohort and 30% of the tiles from the TCGA and GRAMPIAN datasets were used. Domain adversarial training did not involve imCMS class information. Our experiments demonstrate domain adversarial learning is critical to train a classifier that is suitable for this task [**Table 3**].

**Table 3:** Area under the curve (AUC) with 95% confidence intervals achieved by the imCMS classifier trained by domain-adversarial training.

**A) FOCUS**
| **CMS** | **5x (n slides = 506, n patients = 276)** |  |  |  |  |  |
| --- | --- | --- | --- | --- | --- | --- |
|  | **Model 1** | **Model 2** | **Model 3** | **Model 4** | **Model 5** | **Ensemble** |
| **CMS1** | 0.87 (0.78,0.96) | 0.84 (0.73,1.04) | 0.95 (0.91,1) | 0.81 (0.7,0.93) | 0.75 (0.63,0.89) | 0.84 (0.81,0.88) |
| **CMS2** | 0.89 (0.83,0.96) | 0.95 (0.92,1.01) | 0.94 (0.91,0.99) | 0.88 (0.82,0.96) | 0.83 (0.75,0.93) | 0.89 (0.86,0.92) |
| **CMS3** | 0.94 (0.89,0.99) | 0.98 (0.96,1.01) | 0.89 (0.8,1.02) | 0.85 (0.76,1) | 0.94 (0.9,1) | 0.92 (0.89,0.95) |
| **CMS4** | 0.89 (0.83,0.99) | 0.89 (0.82,0.96) | 0.8 (0.69,0.95) | 0.9 (0.84,0.98) | 0.84 (0.77,0.93) | 0.85 (0.81,0.88) |
| **Macro-average** | 0.89 (0.85,0.94) | 0.91 (0.87,0.95) | 0.89 (0.83,0.96) | 0.86 (0.8,0.93) | 0.84 (0.77,0.91) | 0.88 (0.86,0.9) |
| **Micro-average** | 0.88 (0.83,0.92) | 0.92 (0.88,0.97) | 0.91 (0.87,0.95) | 0.88 (0.84,0.93) | 0.83 (0.79,0.87) | 0.88 (0.87,0.91) |

**B) TCGA**
| **CMS** | **5x (n slides = 366, n patients = 365)** |  |  |  |  |  |
| --- | --- | --- | --- | --- | --- | --- |
|  | **Model 1** | **Model 2** | **Model 3** | **Model 4** | **Model 5** | **Ensemble** |
| **CMS1** | 0.84 (0.79,0.91) | 0.83 (0.78,0.87) | 0.81 (0.75,0.88) | 0.81 (0.75,0.89) | 0.81 (0.75,0.86) | 0.84 (0.79,0.89) |
| **CMS2** | 0.79 (0.74,0.84) | 0.81 (0.77,0.87) | 0.79 (0.74,0.85) | 0.8 (0.76,0.86) | 0.79 (0.74,0.85) | 0.83 (0.78,0.87) |
| **CMS3** | 0.82 (0.76,0.89) | 0.75 (0.68,0.83) | 0.8 (0.73,0.87) | 0.81 (0.73,0.9) | 0.82 (0.75,0.9) | 0.83 (0.76,0.9) |
| **CMS4** | 0.73 (0.68,0.8) | 0.77 (0.73,0.83) | 0.74 (0.7,0.8) | 0.78 (0.73,0.84) | 0.72 (0.67,0.78) | 0.78 (0.72,0.83) |
| **Macro-average** | 0.8 (0.76,0.83) | 0.79 (0.76,0.83) | 0.79 (0.75,0.82) | 0.8 (0.77,0.84) | 0.79 (0.75,0.82) | 0.82 (0.79,0.85) |
| **Micro-average** | 0.81 (0.78,0.85) | 0.8 (0.77,0.83) | 0.8 (0.78,0.84) | 0.81 (0.77,0.85) | 0.79 (0.77,0.83) | 0.83 (0.8,0.86) |

C) GRAMPIAN
| CMS | 20x (n slides = 205, n patients = 114) |  |  |  |  |  |
| --- | --- | --- | --- | --- | --- | --- |
|  | Model 1 | Model 2 | Model 3 | Model 4 | Model 5 | Ensemble |
| **CMS1** | 0.81 (0.73,0.9) | 0.56 (0.41,0.72) | 0.74 (0.61,0.85) | 0.77 (0.66,0.89) | 0.85 (0.78,0.96) | 0.85 (0.78,0.91) |
| **CMS2** | 0.83 (0.77,0.89) | 0.75 (0.67,0.83) | 0.74 (0.66,0.8) | 0.7 (0.62,0.78) | 0.79 (0.74,0.85) | 0.8 (0.74,0.85) |
| **CMS3** | 0.89 (0.83,0.96) | 0.8 (0.73,0.89) | 0.8 (0.71,0.89) | 0.8 (0.73,0.87) | 0.82 (0.76,0.88) | 0.86 (0.8,0.93) |
| **CMS4** | 0.91 (0.87,0.96) | 0.84 (0.75,0.93) | 0.86 (0.79,0.93) | 0.9 (0.84,0.97) | 0.91 (0.87,0.98) | 0.92 (0.86,0.99) |
| **Macro-average** | 0.86 (0.82,0.9) | 0.74 (0.68,0.79) | 0.79 (0.73,0.84) | 0.79 (0.74,0.83) | 0.84 (0.81,0.89) | 0.85 (0.82,0.89) |
| **Micro-average** | 0.86 (0.83,0.91) | 0.78 (0.74,0.84) | 0.79 (0.75,0.84) | 0.81 (0.76,0.86) | 0.84 (0.81,0.88) | 0.84 (0.8,0.89) |

#### Adjustment of the imCMS classification probability in the GRAMPIAN cohort

Image tiles containing histological features associated with the imCMS1 class in resection specimens (band like lymphocytic infiltration and mucin) were underrepresented in the rectal biopsies in the GRAMPIAN cohort. This resulted in very few biopsy samples considered as imCMS1 with high confidence [**Table 4**] leading us to adjust the slide-level imCMS classification probabilities. To this end, we trained a RF classifier (*42*) with 100 trees of the maximum depth of 2 with 5-fold cross-validation and only used the results from the test folds to avoid biased adjustment.

**Table 4:** Percentages of image tiles classified as different imCMS classes.

| Prediction | FOCUS<br>*n tiles = 410481* | TCGA<br>*n tiles = 93161* | GRAMPIAN<br>*n tiles = 43754* |
| --- | --- | --- | --- |
| imCMS1 | 25% | 20% | 4% |
| imCMS2 | 31% | 49% | 55% |
| imCMS3 | 11% | 15% | 26% |
| imCMS4 | 33% | 16% | 15% |

#### imCMS classification of the CMS unclassified samples

CMS unclassified samples from all three cohorts were re-classified using the imCMS classification algorithm. To this end, we trained a RF classifier (*42*) on the imCMS classification probabilities of classified samples in the cohort and then applied the learnt classifier to the unclassified samples to assign an imCMS call. Note that for the GRAMPIAN cohort, adjustment of the imCMS prediction probabilities were required as described in the previous section.

### Intratumoural heterogeneity of the imCMS classification

#### Cosine similarity

To evaluate whether the imCMS classification captures the heterogeneity of the transcriptomic CMS classification, we measured the similarity between the imCMS prediction probabilities and their CMS counterpart using cosine similarity, i.e.

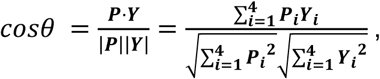

where *P* = [*P*_1_, *P*_2_, *P*_3_, *P*_4_] denotes the imCMS prediction probabilities of a slide, and *Y* = [*Y*_1_, *Y*_2_, *Y*_3_, *Y*_4_] represents the CMS classification probabilities from a RF CMSclassifier (*13*).

#### Assessment of the consistency between the imCMS and CMS classification heterogeneity

We assessed whether the level of similarity between the imCMS prediction probabilities and those of the transcriptomic CMS was better than the level of similarity produced by a random classifier. Samples were stratified according to their primary and secondary CMS profile. For each comparison, a total of 100 random predictions were drawn from a 4-dimensional Dirichlet distribution with a concentration hyperparameter of 1.0 in each dimension in analogy to the imCMS classification probabilities. We calculated the cosine similarities of these random prediction probabilities and the mean of the CMS prediction probabilities.

The median difference between groups was compared using the Wilcoxon rank-sum test and the p-values were adjusted to control false discovery rate (*43*). Any comparison that was highly underpowered due to the sample size (less than 2 data points in one of the populations) was discarded. For each group, outliers were detected via Tukey’s rule (*44*) and removed. To avoid data correlation due to pairs of slides from the same samples, we performed two separate tests in which only one slide from a pair is used in each test. P-values <0.05 were considered statistically significant.

### Survival analyses

Overall survival (OS) in the FOCUS cohort was computed from time of diagnosis of the primary CRC (from 1988 to 2003) until death and was right censored for patients still alive at the date of last known follow-up. OS and data on the progression-free interval (PFI) in TCGA were retrieved from Liu et al (*31*). Patients with less than 1 month of follow-up were excluded. Survival data for FOCUS and TCGA is summarised in [**Tables S6, S7, S8 and S9**]. The GRAMPIAN cohort was not included in the survival analysis due to missing or sparse follow-up data. Univariate Cox proportional hazards analysis was performed to assess the prognostic values of the imCMS classification. Multivariable Cox regression analysis was carried out with TNM stage, age and gender as possible confounding factors following verification of the proportional hazards assumption. P-values <0.05 were considered statistically significant.

### Ethics approval

The use of patient material for cohorts 1 and 3 of the S:CORT program was approved by the ethics commission (REC 15/EE/0241).

## RESULTS

### A deep learning framework for imCMS classification of CRC histology slides

The aim of this study was to develop an image analysis framework to associate features of tissue organisation on standard histology slides with molecular classification and outcome data in CRC patients. Training and test cohorts were selected to represent relevant clinical scenarios in the management of CRC patients including post-operative resection specimens (FOCUS and TCGA) and endoscopic biopsy material (GRAMPIAN). A total of 1,553 slides from three independent datasets were utilised in this study including 666 slides of resection specimens from 362 patients in the FOCUS cohort, 578 slides of resection specimens from 572 patients in the TCGA cohort, and 309 slides from pre-operative biopsies of 175 patients in the GRAMPIAN cohort [**Figure 1A**]. Tumour areas on each slide were annotated by a pathologist and the molecular analysis was performed on material obtained from strict serial sections to derive the CMS calls (*13*) [**Figure 1B**].

The imCMS classifier was trained against CMS calls on the transcriptionally classified samples of the FOCUS cohort and tested on the TCGA and GRAMPIAN cohorts [**Online Methods**]. With the assumption that each CMS class is associated with unique histological patterns localised in different regions of the tumours (*14*), inception V3 deep neural networks (DNN) were trained for prediction of CMS calls for small overlapped image regions (tiles) of 512x512 pixels within the annotated regions [**Figure 1C**]. The size distribution of annotated areas per slide and the number of tiles per slide is shown in [**Figure S1**]. The imCMS class, prediction probability and spatial location for each tile were recorded. An overall imCMS call for each slide was assigned based on the majority classification of tiles [**Figure 1D**].

### imCMS classification is accurate, robust and generalisable

We systematically compared the performance of the imCMS classifier across all three cohorts. For benchmarking against molecular data, all unclassified samples were excluded from the test set. Classification performance was compared using image tiles derived at a) 5x and b) 20x magnification to determine the effect of detail levels. In the FOCUS training cohort, a robust imCMS classification performance of 0.88 AUC (macro-average) was reached [**Tables 1, S2**]. imCMS classification was then tested on the unseen TCGA and GRAMPIAN cohorts [**Tables 1, S2**]. In general, imCMS trained at 5x marginally outperformed classification at 20x on whole tissue sections (AUC FOCUS: 0.88 at 5x vs 0.87 at 20x; TCGA 0.79 at 5x vs 0.78 at 20x), while the 20x imCMS classifier performed better at higher magnification of the endoscopic biopsy specimens (AUC GRAMPIAN: 0.83 at 5x vs 0.85 at 20x). This suggests that training imCMS at higher magnification supports augmentation of morphological features in small tissue samples for imCMS classification. Generalisability was further optimised by adversarial domain training of the imCMS framework, which penalises cohort specific- features during network optimisation [**Online Methods**]. The optimised classifier reached a final classification accuracy of 0.82 AUC on the TCGA cohort and 0.85 AUC on the GRAMPIAN cohort [**Figure 2A and Table 2**]. The correspondence of the CMS and imCMS classification calls for each case is shown in [**Figures 2B, S2**]. Next, we evaluated the consistency of the classification results on pairs of slides obtained from the same patients in the FOCUS and GRAMPIAN datasets. Two H&E slides were generated at different depth levels of each tissue block with at least 4 additional sections cut between for RNA extraction [**Figure S3A**]. Since tissue features at different tissue levels are closely related, a robust classifier would be expected to achieve similar classification results. Indeed, imCMS classification achieved consistent prediction probability between the slide pairs across different CMS classes (Pearson correlation coefficient, FOCUS: 0.89-0.96 and GRAMPIAN: 0.86-0.89, **Figure S3B**).

**Figure 2:**
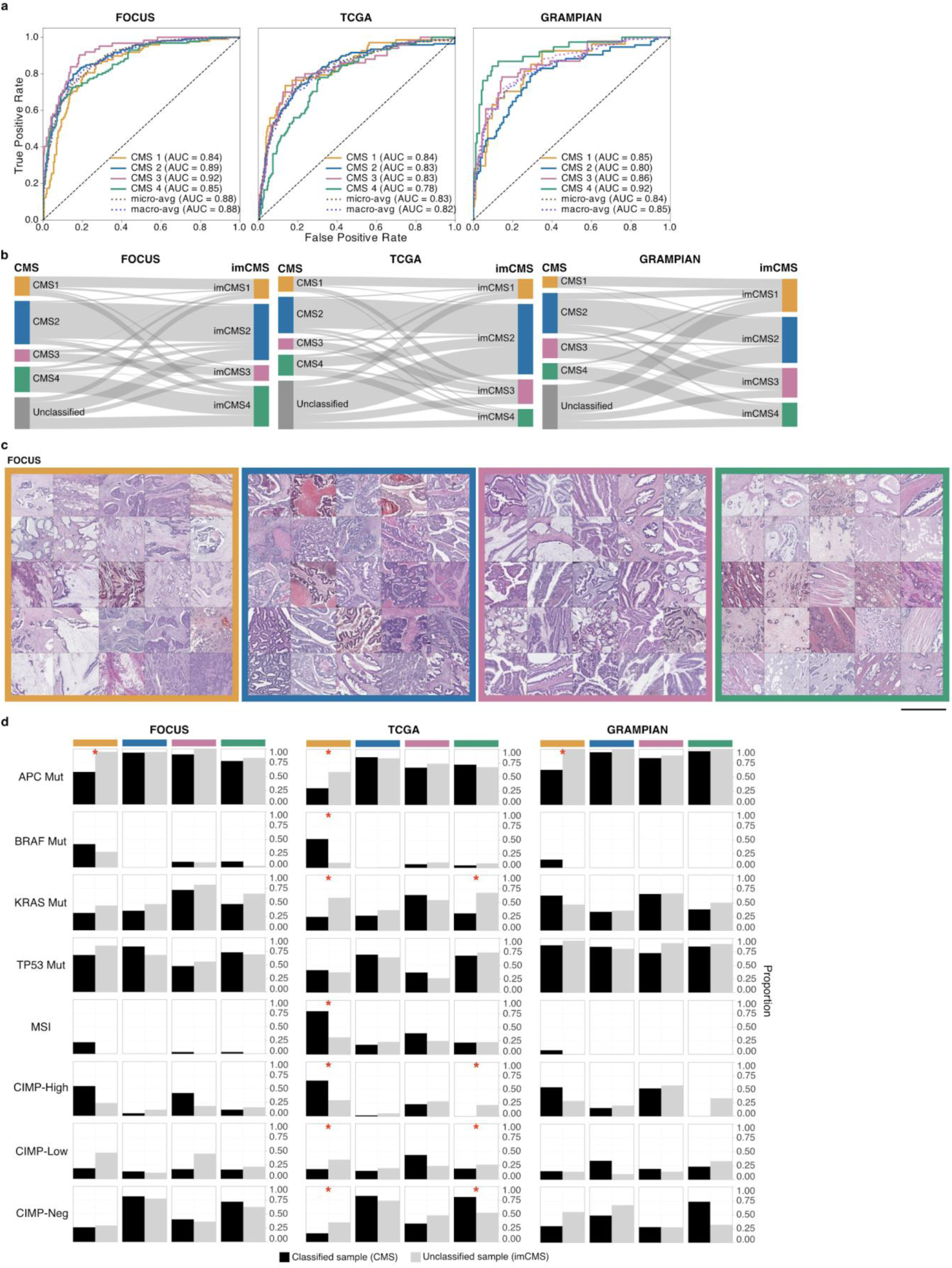
image-based consensus molecular subtype classification. **(A)** Receiver operating curves (ROC) of the imCMS classifier, optimised by the domain adversarial approach, on the FOCUS (n slides = 506, 5x), TCGA (n slides = 366, 5x), and GRAMPIAN cohorts (n slides = 205, 20x). **(B)** Correspondences between CMS and imCMS classes in different datasets. All samples labeled as unclassified by RNA-based CMS calls were successfully re-classified by imCMS **(C)** Examples of image tiles with high prediction confidence for each imCMS class in FOCUS. Histological patterns associated with imCMS 1 are mucin and lymphocytic infiltration. In imCMS2, evident cribriform growth patterns and comedo-like necrosis are observed, while imCMS3 is characterised by ectatic, mucin filled glandular structures in combination with a minor component showing papillary and cribriform morphology. imCMS4 are predominantly associated with infiltrative CRC growth pattern, a prominent desmoplastic stromal reaction and frequent presence of single cell invasion (tumor budding). Scale bar ∼ 1 mm. **(D)** Molecular associations of the CMS classified samples (black) and the CMS unclassified samples that have been classified by imCMS (grey). The molecular profiles of reclassified samples are largely consistent with those of the classified CMS samples. Statistically significant differences (p < 0.05) are marked with a red asterisk.

### Histological patterns associated with imCMS status

To understand which specific morphological patterns associate with imCMS, we extracted and visually reviewed tiles with the highest prediction confidence for each imCMS subtype. The large-scale histology patterns corresponded well with the biological characteristics of the CMS1 and CMS4 classes as predicted from the molecular assay (*13*): Mucinous differentiation and lymphocytic infiltration were associated with imCMS1, and a prominent desmoplastic stromal reaction with imCMS4. imCMS further allowed to visualise and systematically compare the previously poorly defined histological patterns of CMS2 and CMS3 classes. Image tiles associated with high confidence calls of imCMS2 and imCMS3 showed a predominantly glandular differentiation [**Figures 2C, S4A**]. In imCMS2, evident cribriform growth patterns and comedo-like necrosis was observed, while imCMS3 was characterised by ectatic, mucin filled glandular structures in combination with a minor component showing papillary and cribriform morphology. Detailed visualisation of the image representations at the pixel-level corroborated the cellular and tissue components that weigh in on imCMS at high resolution [**Figure S4B**].

### imCMS classification on molecularly unclassified CMS samples

Failure of the transcriptional CMS classification might represent a transition phenotype, intratumoural heterogeneity or might represent technical failure to classify (*13*). We therefore tested the performance of imCMS in samples categorised as unclassifiable by transcriptomic CMS [**Figure 2B**]. As compared to transcriptional classification, imCMS yielded a significantly higher prediction confidence on the molecularly unclassified samples [**Figure S5**]. Successful re-classification is underlined by a direct comparison of the key molecular profiles between classified samples and the imCMS reclassified samples. No major differences between these two groups in the majority of the traits except for CMS1 was found [**Figures 2D, S6, and Table S2**]. However, within the CMS1 subgroup, MSI samples were characterised by higher a priori RF CMS prediction scores (0.69 in MSI+ vs 0.51 in MSI-, p=2x10^−16^, Student’s t-test), leading to a higher probability of accurate identification by CMS. This skewed the proportion of the remaining unclassified samples within the CMS1 subgroup by transcriptional classification towards MSS CRC and explains differences in distribution of MSI- associated molecular features (BRAF, KRAS, CIMP) between the classified and unclassified samples.

### Intratumoural heterogeneity of the imCMS classification

CRC tumours exhibit intratumoural variability in transcriptional features leading to a bias in transcriptional CMS calls introduced by the regions sampled for molecular analysis (*14*). imCMS captures this intrinsic variation in separate predictions for each image tile and provides a model to better reflect and visualise the intratumoural transcriptional heterogeneity of CRC [**Figures 3A, S7a-d**]. We investigated if imCMS heterogeneity was associated with that of the molecular classification. Comparison of the imCMS versus CMS prediction probabilities revealed a high level of agreement between both classification schemes in the majority of the slides [**Figures 3B, S8A**]. We next derived secondary CMS calls from the molecular data [**Figure 3C, Online Methods**] and further looked at the similarity between the corresponding CMS and imCMS prediction probabilities as stratified by primary and secondary CMS calls [**Figure 3D**]. Based on the cosine similarity measure, the match in the variation of the prediction scores was significantly better than by random chance in the majority of groups [**Figures 3D, S8b, Online Methods**], underlining the potential of imCMS to detect and spatially resolve intratumoural heterogeneity in the transcriptional classification of CRC.

**Figure 3:**
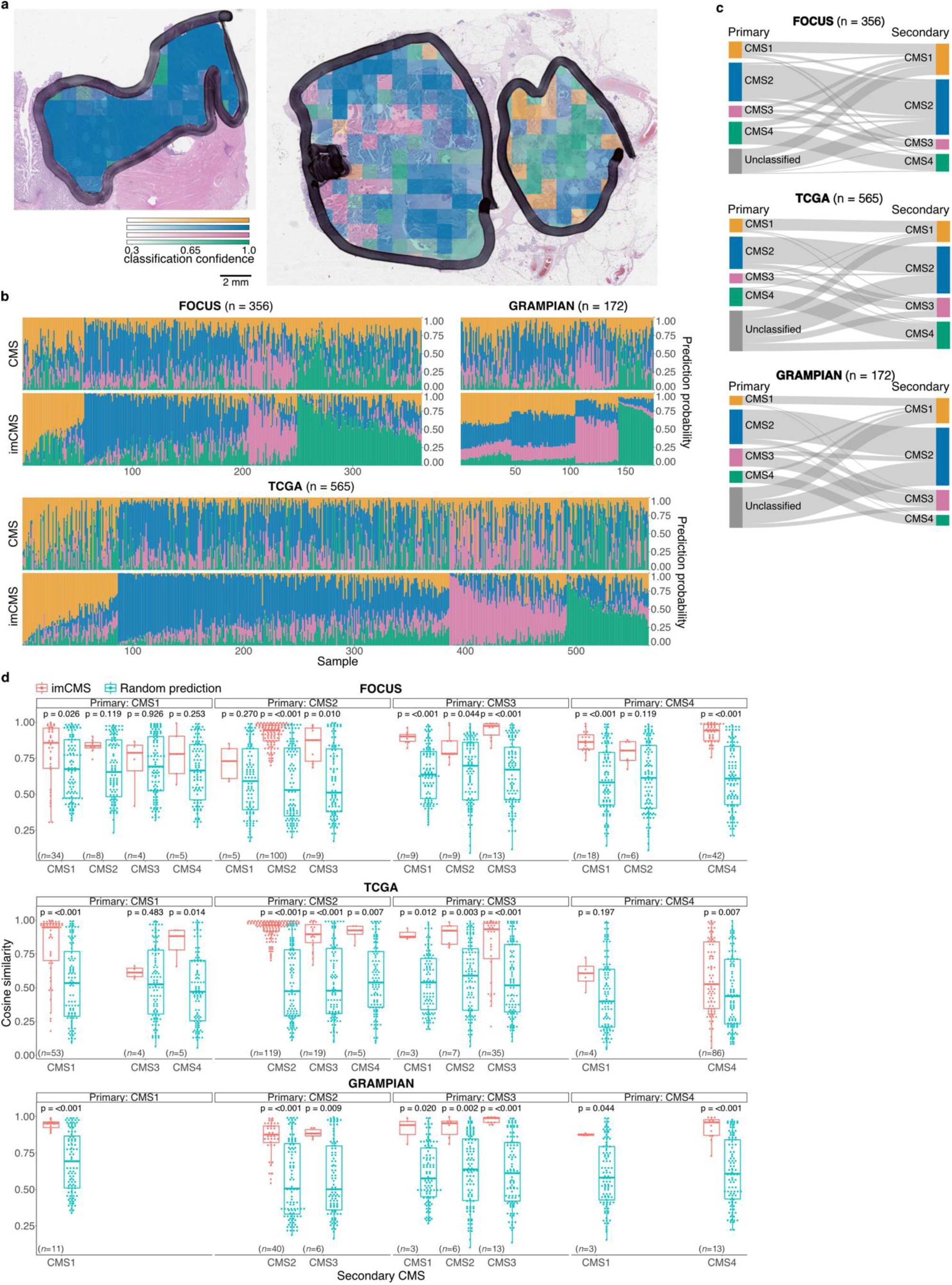
Intratumoural heterogeneity of the imCMS molecular subtypes. **(A)** Visualisation of the regional classification of the imCMS classifier. imCMS classification of a tumour sample can exhibit uniform results (left) or a degree of variation in the predicted imCMS class and the level of confidence (right). The colour overlay indicates the imCMS classes and the opacity reflects the classification confidence. **(B)** Heterogeneity of the CMS and imCMS classification at the slide level. Each bar represents classification probabilities of a sample. **(C)** Heterogeneity of the CMS classification. A secondary CMS call was derived by relaxing the classification threshold of the random forest CMS classifier (*13*). **(D)** Cosine similarity between the imCMS and CMS prediction scores, stratified by the primary and secondary CMS calls. The levels of similarity were compared against those produced by a random classifier. Statistical analysis was performed using Wilcoxon rank-sum test, adjusted for the false discovery rate. P-value < 0.05 was considered statistically significant.

### Prognostic associations by imCMS status

We performed univariate Cox proportional hazard analysis to assess the prognostic value of the imCMS classification as compared to its molecular counterpart. In the FOCUS cohort, patient survival outcomes stratified by imCMS classification were highly in agreement with those of the transcriptional classification [**Figure 4A and Tables S6, S7**]. The prognostic association of the imCMS classification was maintained in multivariate analysis including TNM stage, age and gender, indicating strong potential to stratify risk beyond pathological staging [**Table 5**]. imCMS survival predictions were concordant when the input slides were replaced by sections cut at deeper tissue levels [**Table S3 and Figure S9A**]. For the TCGA cohort, PFI by both imCMS and CMS groups was highly consistent with CMS4 having the poorest prognosis [**Figure 4B and Table S4**]. For OS, the CMS4 group was associated with the worst outcome while imCMS linked the imCMS1 group to adverse outcome [**Figure 4B and Table S4**]. This discrepancy in the TCGA cohort could be explained by a less robust representation of disease biology by OS as compared to PFI but requires additional investigation in subsequent studies. We further explored the application of the imCMS classification for risk stratification in the unclassified samples of the TCGA cohort. In this previously unclassified group, the imCMS4 group was shown to have worse prognosis for both OS and PFI [**Figure S9b and Table S5**].

**Figure 4:**
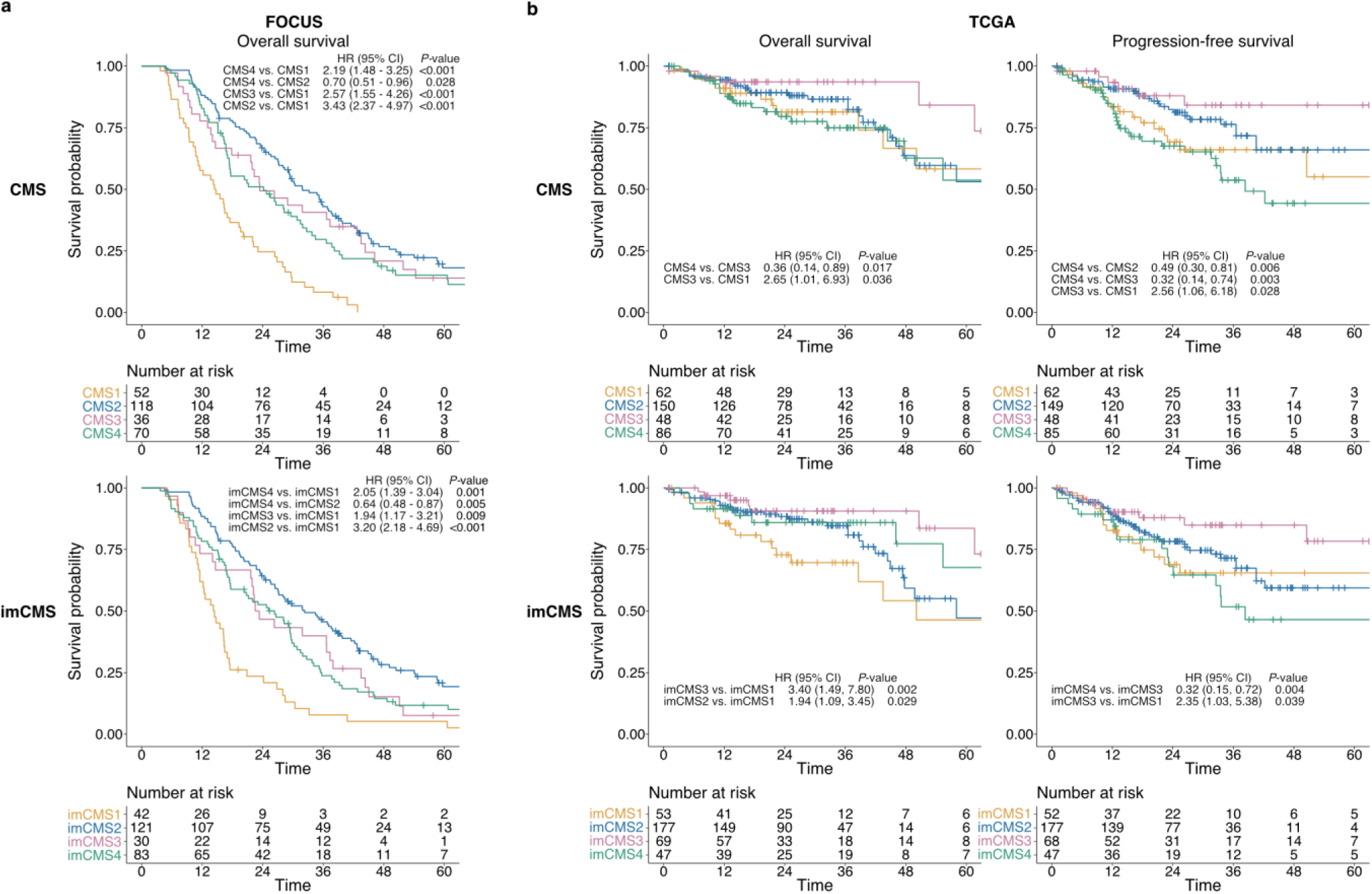
Prognostic associations of the image-based consensus molecular subtypes. Overall survival (OS) and progression-free survival (PFS) outcomes of the **(A)** FOCUS cohort (n=276 cases) and **(B)** TCGA (OS n=346 cases, PFS n = 342 cases) as stratified by the transcriptional-based CMS classification and image-based CMS classification. Kaplan-Meier estimator was used to estimate the survival probability, and pairwise log-rank test and univariate Cox proportional hazards regression were performed between CMS groups and imCMS groups. Hazard ratios (HR) and 95% confidence interval (95% CI) for pairwise comparisons were reported. Test results with p-value < 0.05 were considered statistically significant.

**Table 5:** Multivariate Cox proportional hazards regression on classified samples of the FOCUS cohort.

| FOCUS (n patients=263) | Multivariate survival analysis (adjusted by gender, age, T, N, M) |  |  |  |
| --- | --- | --- | --- | --- |
| Variable | HR | 95%CI Low | 95%CI High | p-value |
| *CMS1 vs CMS2* | **2.60** | **1.68** | **4.02** | **1.72E-05** |
| *CMS3 vs CMS2* | 0.98 | 0.62 | 1.54 | 9.18E-01 |
| *CMS4 vs CMS2* | 1.34 | 0.93 | 1.91 | 1.12E-01 |
| *imCMS1 vs imCMS2* | **2.20** | **1.37** | **3.54** | **1.13E-03** |
| *imCMS3 vs imCMS2* | 1.37 | 0.86 | 2.17 | 1.89E-01 |
| *imCMS4 vs imCMS2* | **1.48** | **1.05** | **2.08** | **2.68E-02** |

## DISCUSSION

H&E slides are generated as part of the standard work-up of any CRC treated by surgical resection (*16, 17*). In the assessment of this histologic material, pathologists are presently limited to the strictly defined set of morphologic and anatomic criteria (*16, 17*). This information supports the definition of broad prognostic risk groups but has no predictive value (*16*). The integration of genomic technologies in the clinical care of CRC patients has immense potential to drive personalised treatment but requires substantial financial, personnel and infrastructure resources (*18*). Combining morphological and molecular pathology to identify genotype-phenotype correlations is a promising approach to extend the amount of clinically relevant information that can be extracted from standard histologic slides (*8*). In this study, we leverage artificial intelligence and image analysis technologies for the development of an image-based taxonomy of CRC with clear biological interpretability and clinical impact. Due to general applicability and low costs, morphomolecular classification of histopathology slides could become a new standard for patient stratification in clinical practice.

We trained and tested our image-based approach towards consensus molecular subtyping (imCMS) of CRC on three independent and well-characterised patient cohorts with availability of digital slides and transcriptional information from the CRUK MRC S:CORT program and TCGA. We specifically focused on relevant clinical scenarios in the management of CRC patients and investigated the imCMS classification of both preoperative biopsies and resection specimens. Our analyses demonstrate that the imCMS classifier is able to predict the consensus molecular signatures of CRC from histological slides with very high accuracy. While tissue features captured at low magnification proved most informative on CRC resection specimens, imCMS could be efficiently adapted for morpho-molecular classification of rectal cancer biopsy fragments at high magnification. Small biopsy fragments have previously proven difficult to analyse using genomic technologies due to the limited amount of tissue available (*19*). Pathologist assessment is therefore usually restricted to the diagnosis of cancer, a select panel of immunohistochemical studies and a limited assessment of additional prognostic features (*17, 20*). Clinically approved assays that are predictive of therapeutic response from biopsy material are presently lacking, with up to 25% of rectal cancer patients gaining no benefit from current radiotherapy and chemotherapy protocols (*21*). As a stemlike (CMS4) transcriptional profile of CRC has been linked to poor prognosis and therapeutic resistance, imCMS could allow for more effective stratification of patients for primary surgery or neoadjuvant treatment (*22, 23*). Prospective studies are warranted to investigate the application of imCMS as a novel clinical stratification tool.

Our analysis demonstrates the feasibility of imCMS classification of both primary colon and rectal resection specimens in the FOCUS and TCGA cohorts. imCMS calls closely matched transcriptional classification for survival stratification, underlining the strong potential of imCMS for translation into the clinical routine. imCMS classification of surgically treated primary CRC could aid pathologists in the identification of aggressive disease for intensified follow-up and chemotherapy trials (*1*). In advanced disease, the development of molecular stratifiers for the prediction of treatment response is of critical importance to balance care and overtreatment. No clinically approved tests are currently available to predict chemotherapy response in metastatic CRC with as many as 20 patients statistically needed to receive the combination treatment with 5-Fluorouracil and Oxaliplatin to achieve long term (>3 year) disease free survival for one individual (*22*). Beneficial effects are set off by considerable toxicity including debilitating chronic peripheral neuropathy in up to 50% of cases (*24*). Transcriptional classification of CRC has shown promise to stratify survival outcomes and response to treatment in retrospective analyses but requires further validation (*22, 23*). imCMS represents a readily translatable and cost-effective approach for further investigation of treatment outcomes in existing retrospective cohorts and future clinical trials.

Limited generalisability of image analysis algorithms is a well-recognised problem in the setting of limited training sets and poorly annotated ground truth data (*25*). We addressed the problem of sample diversity by training the imCMS classifier on histological samples sourced from multiple institutes (n=59) participating in the FOCUS trial. Domain adversarial training was used to minimise the classification weight of cohort dependent features in the final models (*26*). The ensemble of multiple models, analogous to consensus of experts’ opinions reduces the bias of individual predictions (*27*). High-level annotations were guaranteed by a strict protocol where each H&E section used for digital image analysis was followed by slides cut for molecular profiling with precisely matched annotations. This allowed us to directly associate transcriptional signatures with histological phenotypes in CRC at unprecedented resolution. RNA expression signatures represent both tumour intrinsic and microenvironment related signals which are intimately linked to CRC phenotypes with distinct biological characteristics and disease outcomes (*1, 13*). imCMS highlighted the well-known morpho-molecular associations with inflammatory infiltrates (imCMS1) and a prominent stromal reaction (imCMS4) but also identified novel morphological features in association with high-confidence calls of imCMS2 and imCMS3 while robustly reproducing the known molecular associations of transcriptionally derived CMS subtypes. Our study underlines that convolutional neural networks excel in their ability to learn relationships of tissue compartments as a whole and to identify relevant patterns with clear morphological interpretability.

Transcriptomic CMS was released as the most robust molecular classification in CRC and the basis for clinical stratification and targeted intervention (*1, 13*). However, some key issues hamper clinical implementation of CMS such as the inability to obtain reliable calls from single samples. Two methods to call CMS were released by the original authors based on RF and single sample prediction (*13*). The former provides reliable classification but is cohort-dependent and requires a high minimum number of samples while the latter generates calls on single samples with limited quality leading to underutilisation. Another problem is that some samples do not show enough evidence to make calls by either method leading to a substantial number of cases left as unclassified. Inconsistent classification calls could also be an expression of intratumoural heterogeneity or representative of a transition phenotype which is of considerable biological interest (*1, 13*). Spatial heterogeneity is an additional confounder that can result in CMS misclassification (*14*). imCMS is able to overcome all these problems. imCMS calls are intrinsically generated for single samples. Notably, imCMS images visualise heterogeneity through tile-based classification calls with a cell size of 512 × 512 pixels, allowing us to derive quantitative prediction scores with biological interpretability. Here, we show that transcriptionally unclassified samples tend to have higher heterogeneity of the image-based classification results as compared to the CMS classified samples. Importantly, all CMS unclassified samples were successfully reclassified by imCMS and their molecular characteristics as well as survival profiles closely resembled those classified by sequencing methods. These results suggest that imCMS performs reliably in samples categorised as unclassified by transcriptional profiling and indicates that different molecular profiles within CMS subgroups may be biological rather than technical. Re-classification by imCMS achieved significantly higher confidence for sample categorisation than transcriptional profiling. To further investigate sample heterogeneity, we bioinformatically derived secondary CMS calls from all samples and investigated the similarity of the CMS and imCMS prediction probabilities for primary and secondary calls. imCMS captured secondary calls with high accuracy based on a cosine similarity measure between transcriptional and image-based classification. Taken together, imCMS allows for the first time to localise sources of heterogeneity on the original tissue slide and to understand, control and further investigate sources of heterogeneity in the transcriptional classification of CRC. In addition, imCMS is a versatile tool to address deficiencies in transcriptional profiling that may arise due to low amounts or quality of RNA, an expected problem in clinical FFPE blocks.

With this paper we demonstrate that it is possible to identify CMS on the basis of tissue morphology. The possibility of identifying morphological correlates that are associated with molecular subtypes opens new opportunities for in vitro diagnostics. However, the application of image-based patient stratification is presently limited by the availability of digital pathology infrastructure in routine diagnostic practice. This is met by broad scale initiatives for digitalization of medical infrastructure on a national and international level (*28, 29*). Centralized testing could further compensate for the availability of computing infrastructure in low resource settings. Prospective validation of imCMS in independent studies will be critical to clinical translation. This includes both applications as a tool that could rationalize which cases would need confirmatory testing as well as stand-alone testing in cases where genomic methods fail to provide reliable classification. We hypothesise that the general principle can be applied not only to other cancer types but also to other diseases. It will therefore lay the foundation of a more systematic integration of image-based morphological analysis and molecular stratification.

## ACKNOWLEDGMENTS

The S:CORT consortium is an Medical Research Council stratified medicine consortium jointly funded by the MRC and CRUK. This work was further supported by the National Institute for Health Research (NIHR) Oxford Biomedical Research Centre. Computation used the Oxford Biomedical Research Computing (BMRC) facility, a joint development between the Wellcome Centre for Human Genetics and the Big Data Institute supported by Health Data Research UK and the NIHR Oxford Biomedical Research Centre. J. Rittscher is supported through the EPSRC funded Seebibyte programme (EP/M013774/1). VHK gratefully acknowledges funding by the Swiss National Science Foundation (P2SKP3_168322/1 and P2SKP3_168322/2), the Werner-Hedy Berger Janser Foundation and the Promedica Foundation. The authors thank Aurelien de Reynies for advice on CMS calling in FFPE blocks, Claire Butler and Michael Youdell for excellent managing in S:CORT and the MRC Clinical Trials Unit who provided the clinical data from the FOCUS trial with permission from the FOCUS trial steering group. We would further like to thank Indica Labs for providing the HALO^TM^ software. The results published or shown here based in part upon data generated by the TCGA Research Network: http://cancergenome.nih.gov/ established by the NCI and NHGRI. Information about TCGA and the investigators and institutions who constitute the TCGA research network can be found at http://cancergenome.nih.gov. We would specifically like to thank all patients who have consented to take part in S:CORT and TCGA. The views expressed are those of the author(s) and not necessarily those of the NHS, the NIHR or the Department of Health.

## DATA AVAILABILITY STATEMENT

The datasets generated during and/or analysed during the current study are available from the corresponding authors on reasonable request.

## AUTHORS CONTRIBUTIONS

TM, JR, IT and VHK jointly conceived the study. KS, ED, TM, JR and VHK designed the study; KS, ED, TM, JR, VHK drafted the manuscript; KS, ED, SR, KR, ABl, AC, CH, CW, IT, ABe, UMcD, PD, SW, GIM, LMS, MS, PQ, TM, VHK obtained and categorised clinicopathological and molecular data. KS, ED, TM, JR, VHK performed data interpretation. CV and SL provided important intellectual input, provided critical resources or funding, and critically reviewed the study design. KS, ED, ABl, C-HW performed bioinformatic and statistical analysis. All authors have read and given approval of the final manuscript.

## Supplementary Figures S1-S9

**Figure S1:**
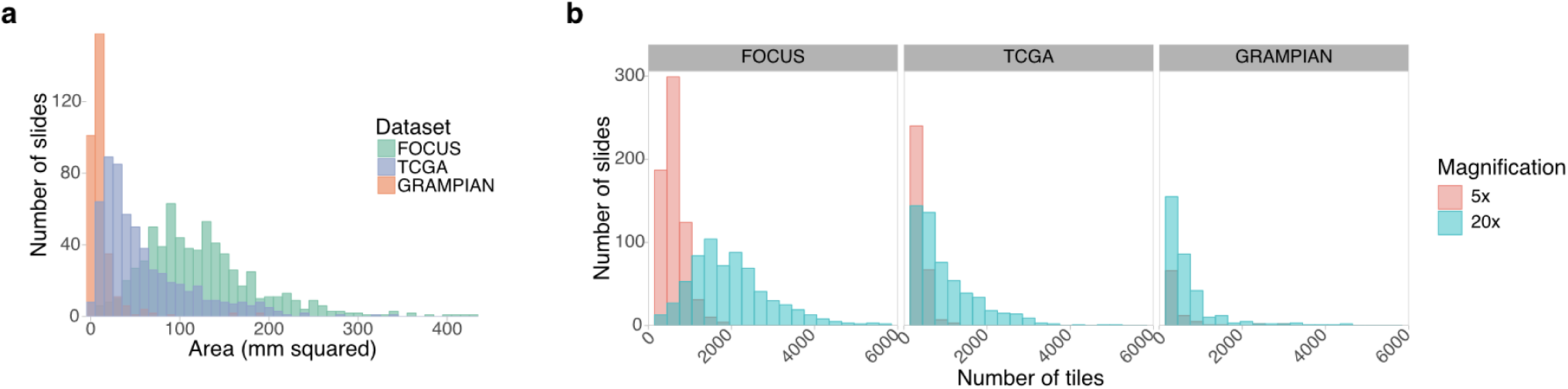
Slide statistics. **(A)** The distribution of annotated tumour areas in different datasets. **(E)** The distribution of the number of tiles extracted from the annotated regions at 5x and 20x.

**Figure S2:**
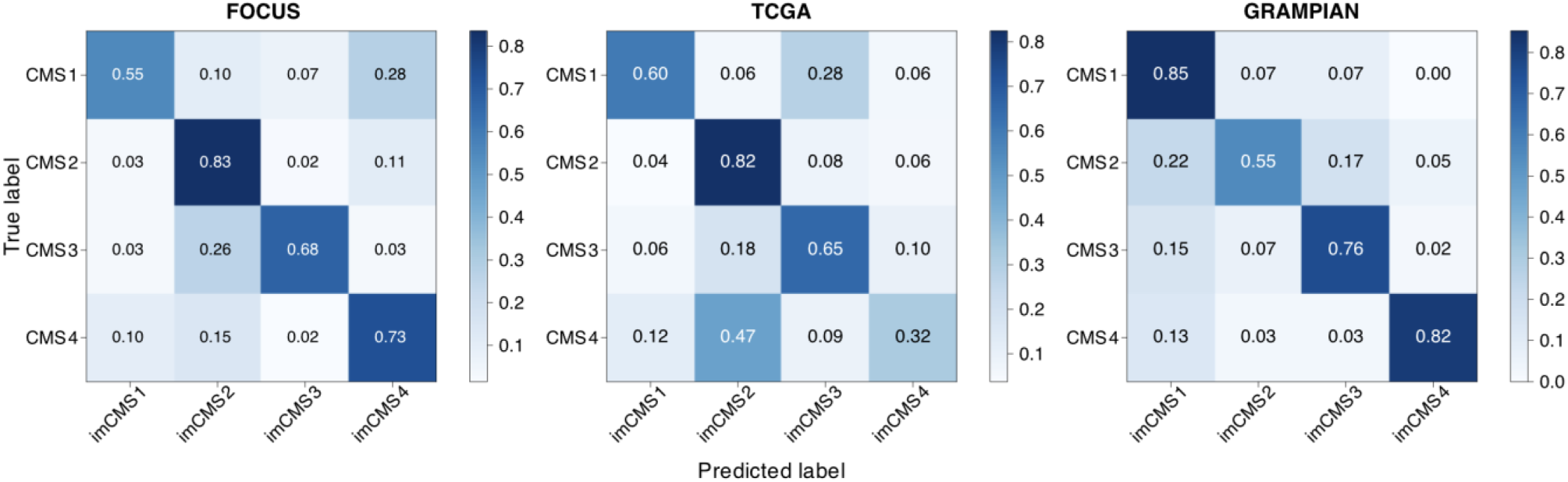
imCMS classification. Confusion matrices showing the classification performance of the imCMS model on different datasets. A sample is assigned to the imCMS class with the maximum prediction score (i.e. highest number of tiles in the slide).

**Figure S3:**
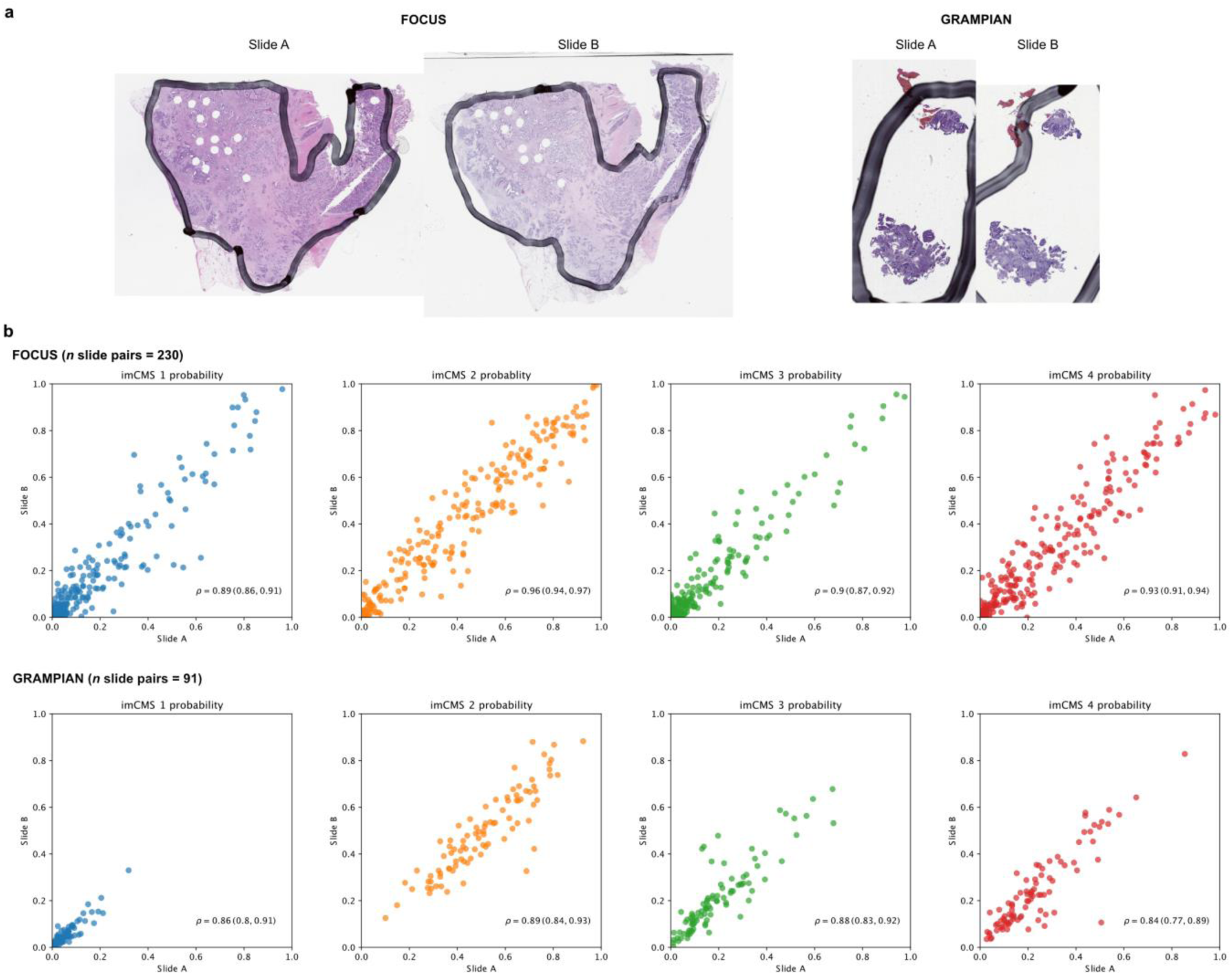
Consistency of the prediction probability. **(A)** Examples of pairs of slides from the FOCUS and GRAMPIAN datasets. **(B)** Pearson correlation coefficient of the predicted probabilities between pairs of slides.

**Figure S4.**
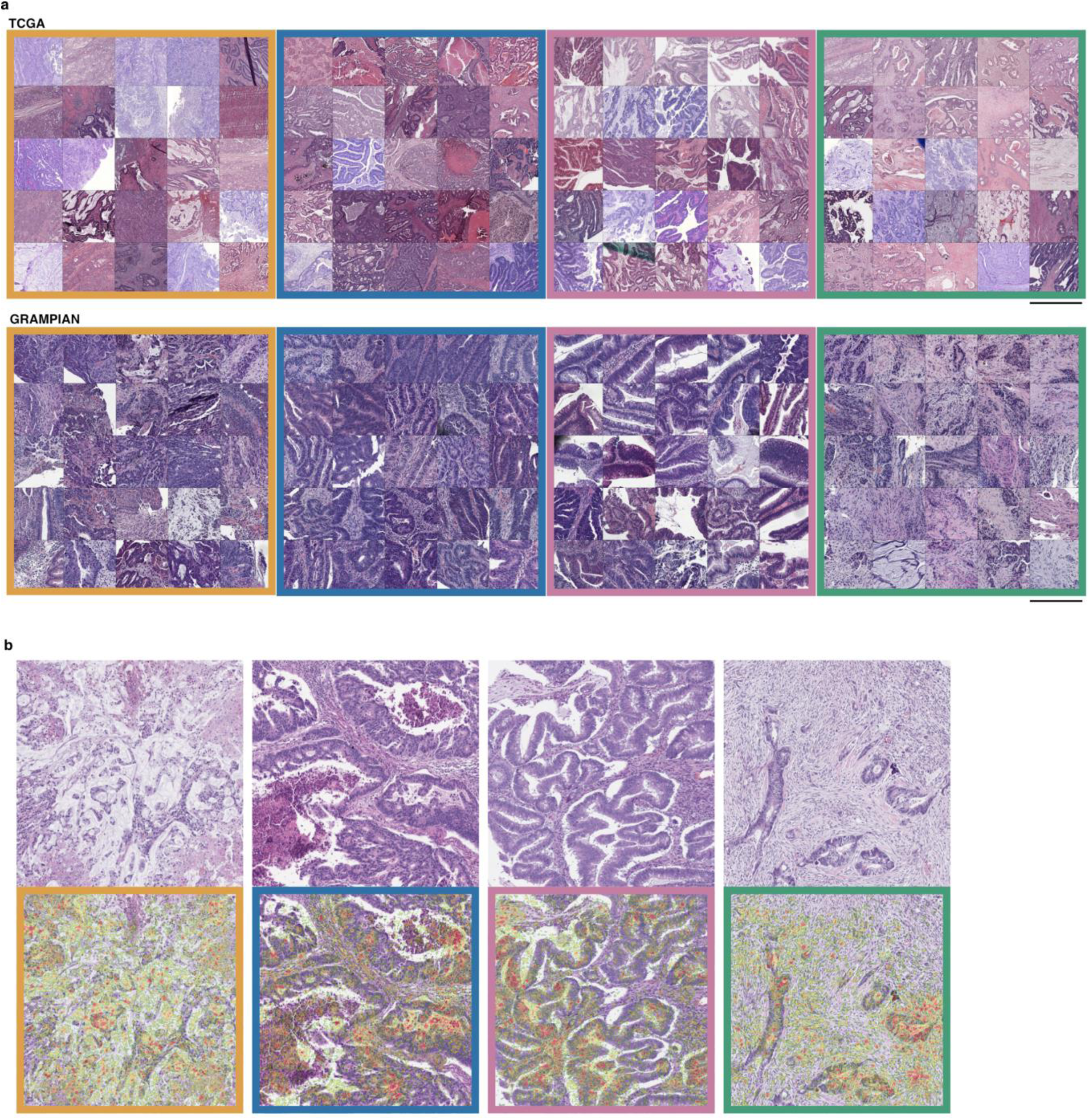
Morphological correlates of the imCMS classes. **(A)** Example image tiles with high prediction confidence from the TCGA cohort (scale bar ∼ 1 mm) and the GRAMPIAN cohort (scale bar ∼ 255 microns). **(B)** Pixel locations important for the class decision are highlighted. The order of importance is represented as a gradient between green and red, where red indicates the highest level of importance. The highlighted pixel locations correspond largely to lymphocyte and mucin in imCMS1, tumour areas in imCMS2 and imCMS3, and infiltrative tumour front and stroma in imCMS4.

**Figure S5:**
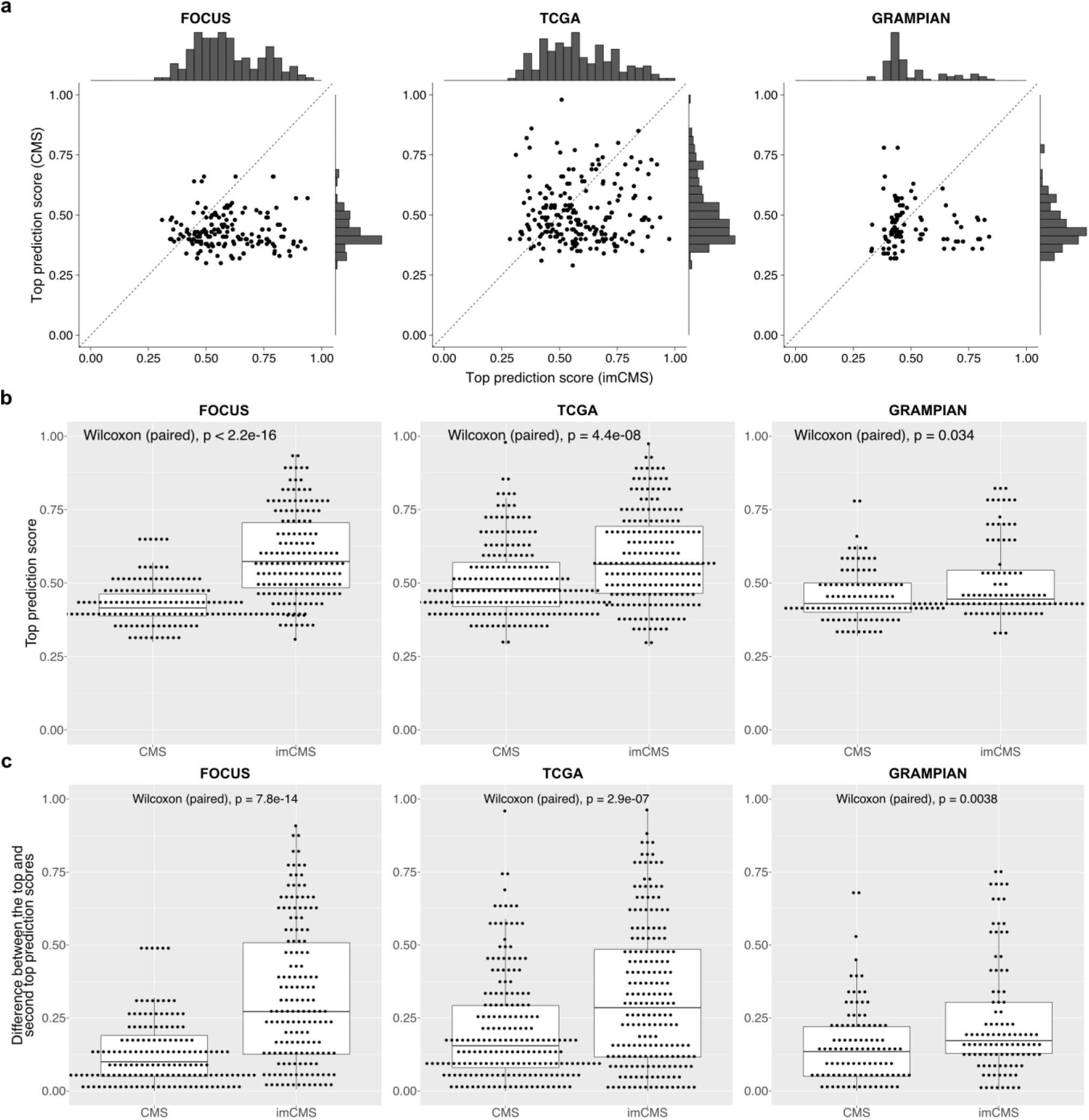
Comparison of the prediction confidences of the CMS and imCMS classifiers in the CMS unclassified samples. **(A)** Correspondences between the top CMS and imCMS prediction scores. **(B)** The top imCMS prediction scores are significantly higher than the corresponding CMS prediction scores in all datasets (Wilcoxon signed rank test, p-values < 0.05). **(C)** The differences between the top and the second top prediction scores produced by the imCMS classifier are significantly larger their CMS counterparts (Wilcoxon signed rank test, p-values < 0.05).

**Figure S6:**
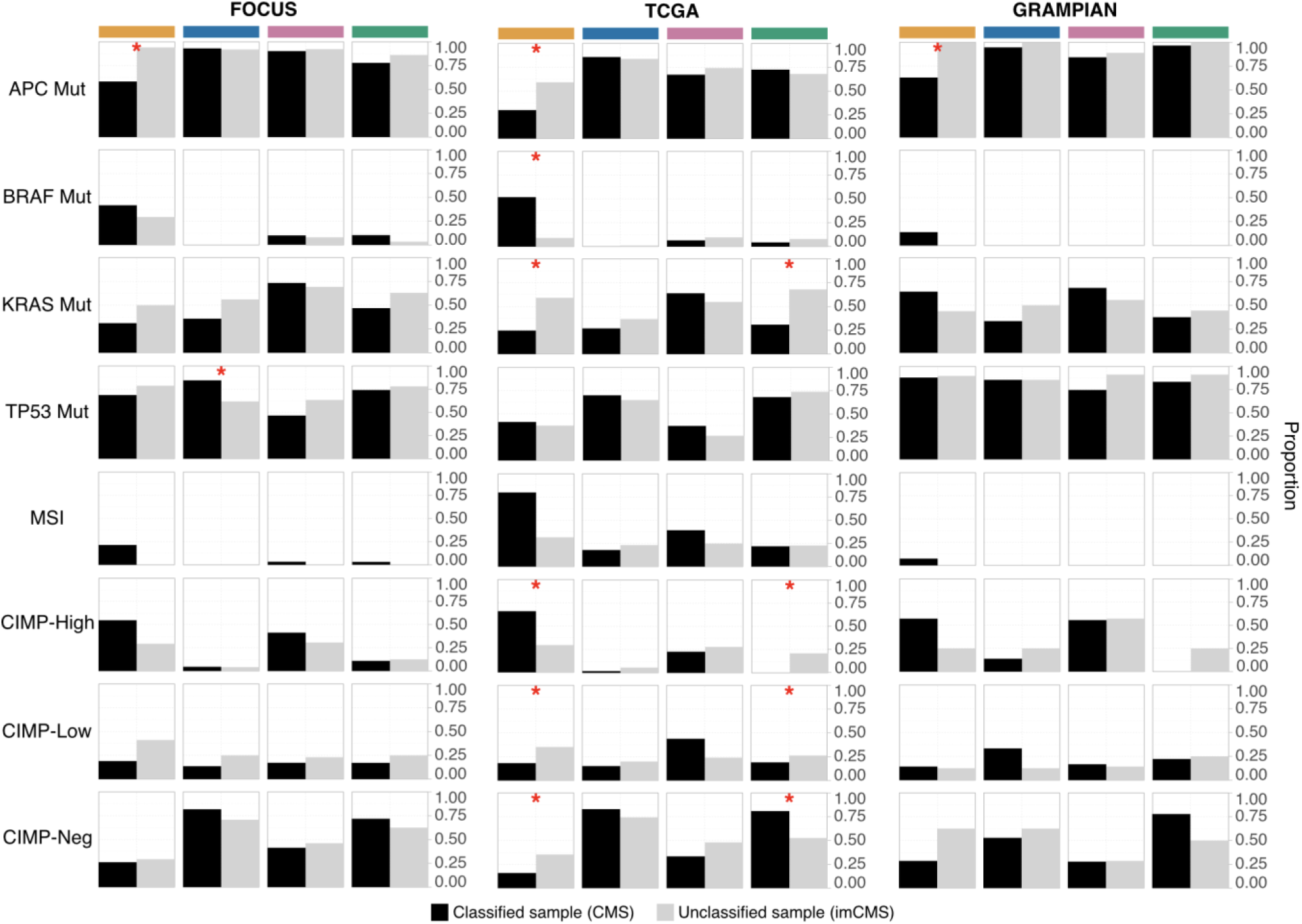
imCMS classification of the CMS unclassified samples. Molecular associations based on the 2nd slide of the CMS classified samples (black) and the CMS unclassified samples that have been classified by imCMS (grey). A significantly different profile (p < 0.05) is marked with a red asterisk.

**Figure S7:**
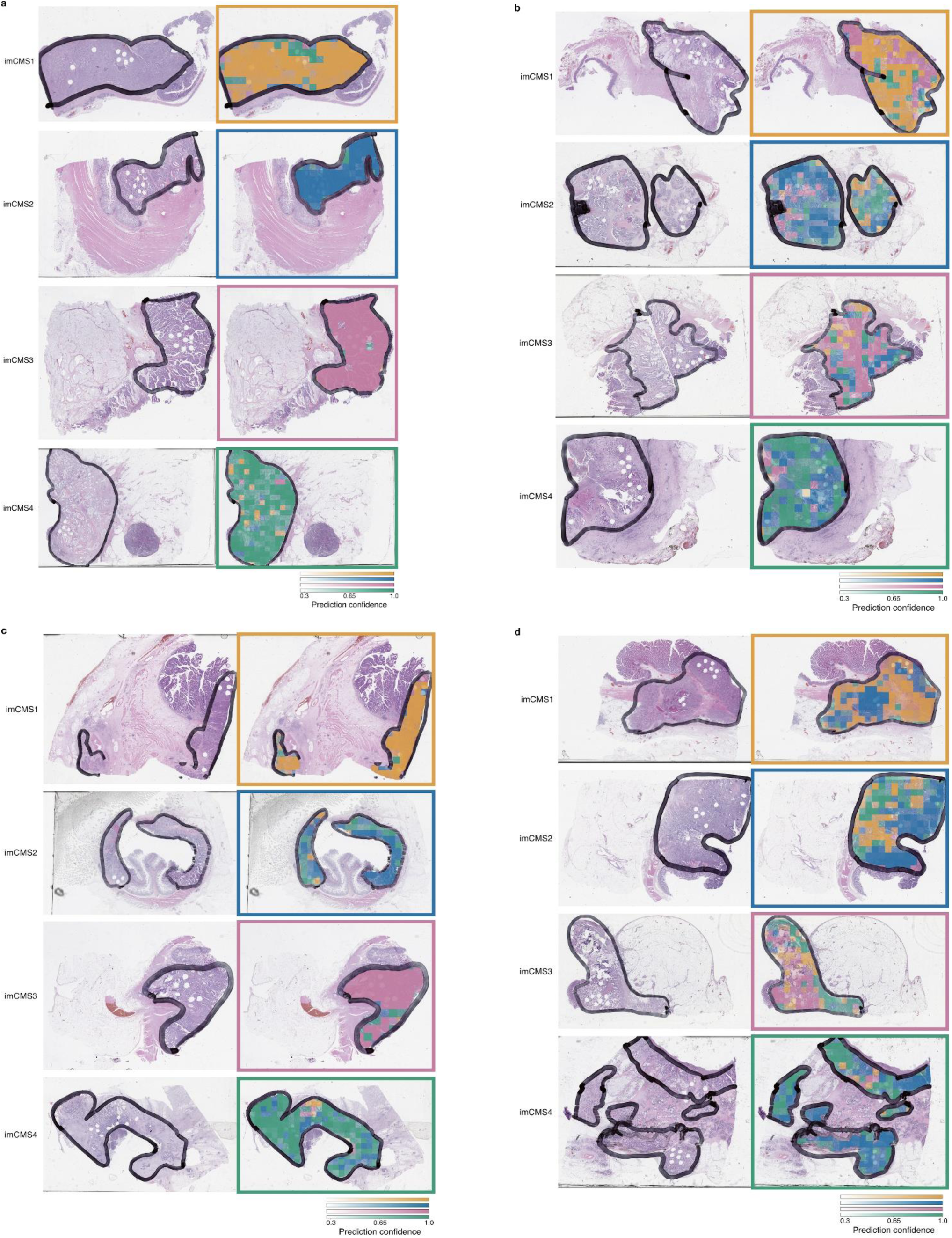
Intratumoural heterogeneity of the imCMS prediction. The heterogeneity of the imCMS prediction per slide can be observed both in the form of the variation in the predicted classes and the variation in the levels of the prediction confidence. **(A)** CMS classified samples with a low level of imCMS prediction heterogeneity. **(B)** CMS classified samples with a high level of imCMS prediction heterogeneity. **(C)** CMS unclassified samples with a low level of imCMS prediction heterogeneity. **(D)** CMS unclassified samples with a high level of imCMS prediction heterogeneity.

**Figure S8:**
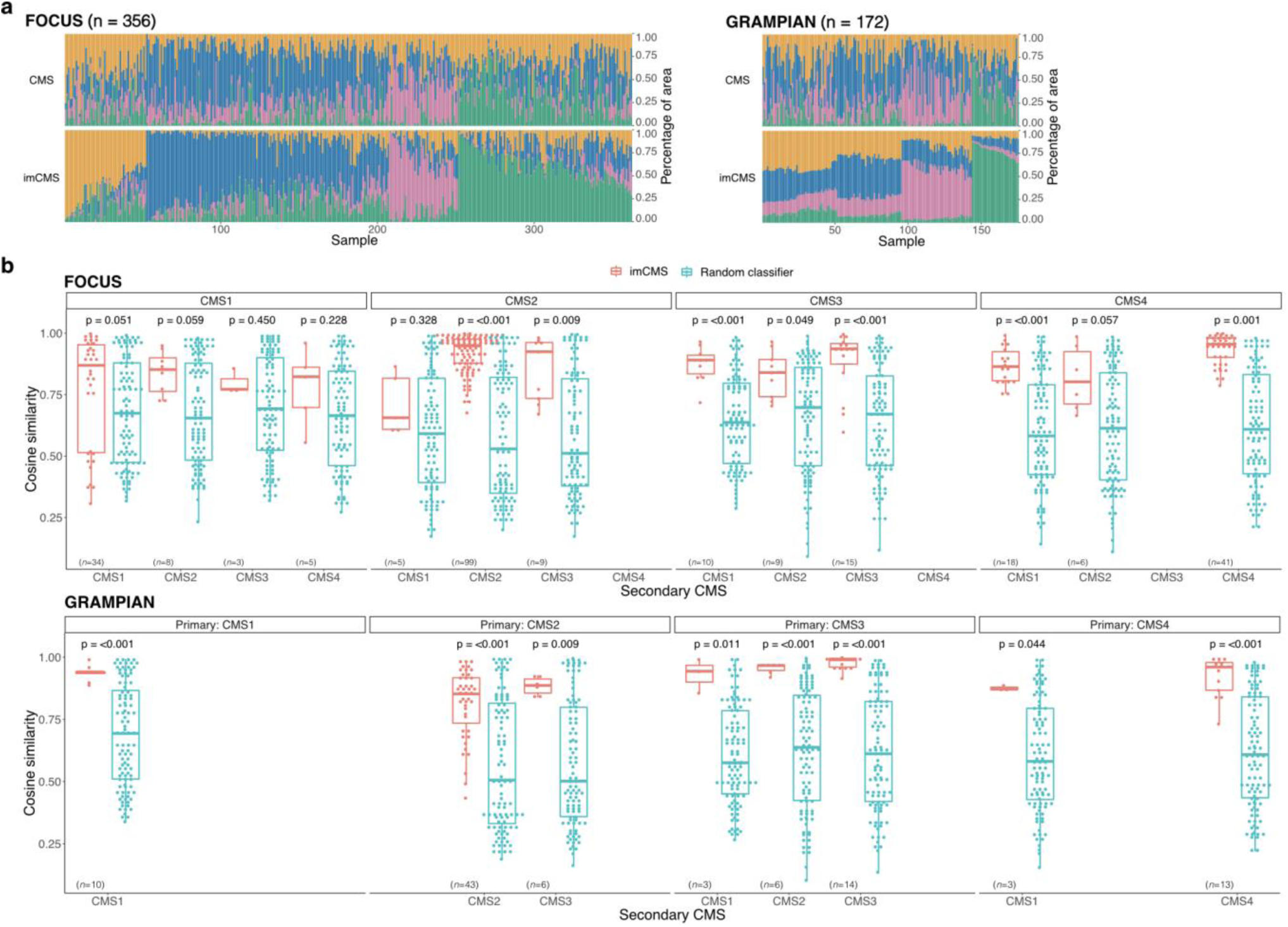
Intratumoural heterogeneity of the imCMS prediction (2nd slides). **(A)** Heterogeneity of the CMS and imCMS classifications. Each bar represents classification probabilities of a sample. **(B)** Cosine similarity between the imCMS and CMS prediction scores, stratified by the primary and the secondary CMS calls. The levels of similarity were compared against those produced by a random classifier. Statistical analysis was performed using Wilcoxon rank-sum test, adjusted for the false discovery rate. P-value < 0.05 was considered statistically significant.

**Figure S9:**
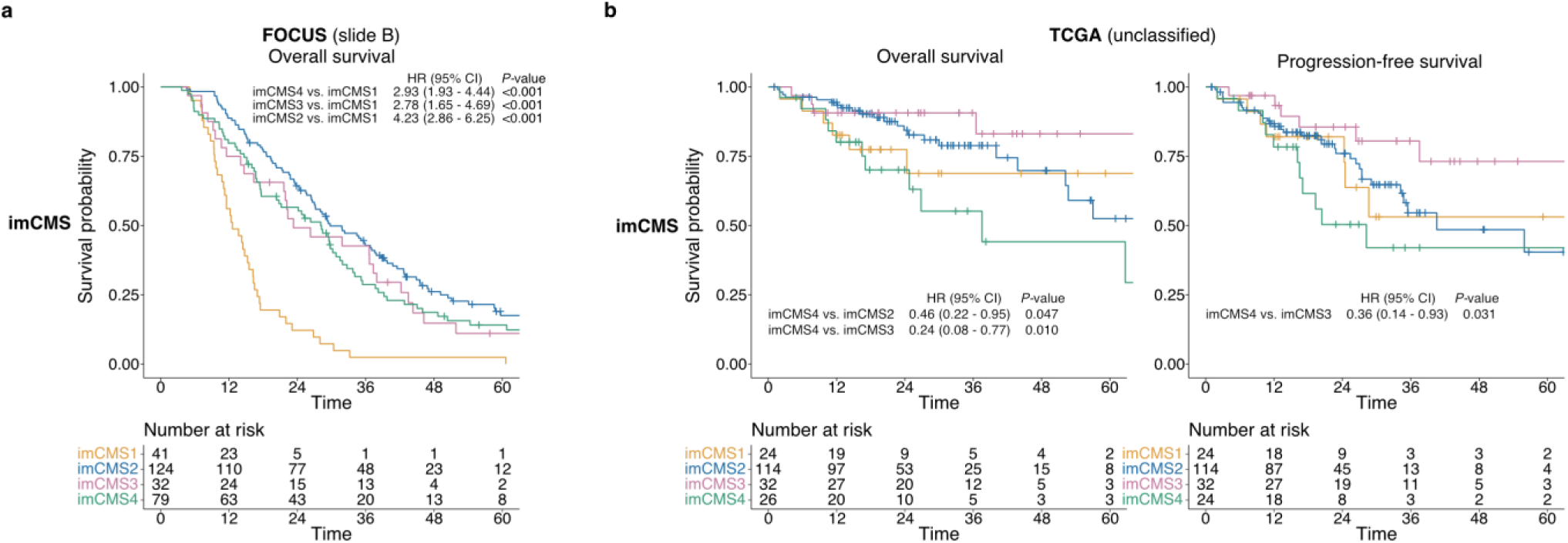
Prognostic associations of the imCMS classification. **(A)** Overall survival analysis based on the 2nd slides of the FOCUS cohort (n=276). **(B)** survival outcomes of the unclassified samples (n=196 cases) from TCGA cohort as stratified by imCMS classification.

## Supplementary Tables S1-S5

*Please see separate datafiles*

**Table S1: Clinicopathological and molecular associations of the datasets (FOCUS, TCGA, GRAMPIAN)**

**Table S2: Molecular associations of CMS classified samples versus CMS unclassified samples (reclassified by the imCMS classification)**

**Table S3: Univariate Cox proportional hazards regression on classified samples of the FOCUS cohort**

**Table S4: Univariate Cox proportional hazards regression on classified samples of the TCGA cohort**

**Table S5: Univariate Cox proportional hazards regression on unclassified samples of the TCGA cohort**

